# Theory Note: Emergence of a Proportional-Derivative Control Law from Two Coupled Oscillating Brain Circuits Near Synchrony

**DOI:** 10.64898/2026.07.13.738181

**Authors:** Omar Refy

## Abstract

Oscillations and oscillatory synchronization are pervasive in motor circuits, where their role in rhythm generation and entrainment is well established but their role in feedback control of movement remains unclear. Here I show analytically that two oscillators of any type, coupled through a delayed interaction that is an odd function of their phase difference, necessarily implement a proportional-derivative (PD) control law in the near-synchrony limit. The proportional gain follows from the slope of the coupling function and the derivative gain is set by the coupling delay, so that PD control emerges with no additional machinery. Simulations confirm that such oscillators reproduce ideal PD step responses near synchrony and that control quality degrades systematically away from it. This establishes a direct, model-independent bridge between oscillatory synchronization and feedback control, and suggests concrete experimental signatures for candidate systems.

## Introduction

Oscillatory activity is a hallmark of the motor system. Rhythms appear in cortical motor areas, in corticomuscular and corticospinal coherence, in central pattern generators, and in the synchronization between motor cortex and the spinal motoneuron pool during sustained contraction [1–4]. The functional roles usually attributed to these rhythms are rhythm generation, temporal coordination, entrainment, and inter-areal communication through coherence [2,4,5]. What such oscillations contribute to the moment-to-moment *feedback control* of movement (i.e. the regulation of limb state against perturbations) is far less clear.

Synchronization of coupled oscillators is itself a mature subject. The dynamics of phase-coupled oscillators, including the canonical case of odd (sinusoidal) coupling in the Kuramoto model and its delayed variants, have been studied in great depth [6–11]. Coupled-oscillator descriptions of biological rhythms are likewise not new [3,4,5]. These treatments overwhelmingly emphasize locking, stability, and collective modes, while feedback control is treated as a separate principle invoked to explain how movement is regulated and how perturbations are rejected [12,13].

The results presented below connect the two. Here, I show that two near-synchronous oscillators interacting through delayed coupling, odd in their phase difference, necessarily implement a PD control law on the plant they drive. In this formulation, synchronization dynamics serve as the implementation of feedback control.

### PD control emerges near synchrony

Consider two oscillators (Figure 1A): a lead oscillator that encodes a desired plant state and a follower oscillator that drives a plant with state *x*(*t*).

**Figure 1.**
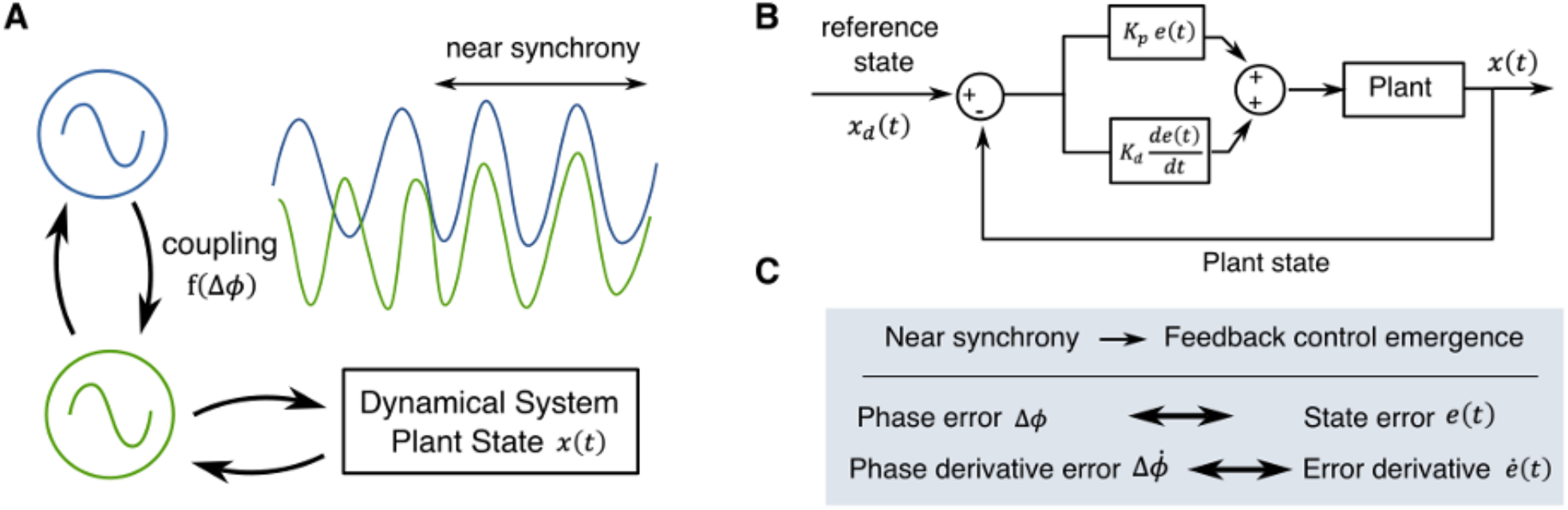
Emergence of PD-like control from two coupled oscillators near synchrony. (A) Two coupled oscillators driving a plant; the follower oscillator’s phase tracks the plant state. (B) Block diagram of an equivalent PD controller. (C) Mapping from phase-difference dynamics to PD control terms: near synchrony the phase error maps to the state error and the phase-derivative error to the error derivative.

#### Assumptions

i. The lead phase encodes the target and the follower is tightly coupled to the plant on a fast time scale, so its phase encodes the plant state:

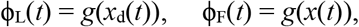

where the encoding map *g* is, for simplicity, shared by both oscillators. We further take *g* to be locally linear near the operating point, with constant slope *g*′, so that *g*′ relates phase increments to state increments, δϕ = *g*′ δ*x*.
ii. The coupling Γ is an odd function of the phase difference Δϕ = ϕ_L_ − ϕ_F_:

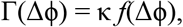

with *f* odd, so the follower is advanced when lagging and retarded when leading [9,14,15]. Kuramoto-type synchronization theory uses a sine coupling, which falls in this class.
iii. The interaction carries a fixed delay δ on the reference (lead) pathway, large compared with the follower-to-plant delay, so the coupling at time *t* compares the delayed lead phase with the present follower phase: Γ(*t*) = κ *f* [ϕ_L_(*t*−δ) − ϕ_F_(*t*)], where *f* is an odd function.

#### Derivation

Because *f* is odd, linearizing near synchrony (Δϕ ≈ 0) replaces *f* by its argument. Expanding the delayed lead phase to first order, 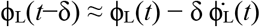, and using 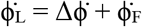, the argument becomes 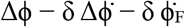, so

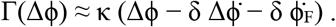

Substituting the encodings from (i) with the error *e*(*t*) = *x*_d_(*t*) − *x*(*t*) and 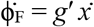 gives

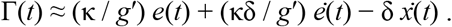

This is a PD control law together with an additional damping term (Figure 1B,C):

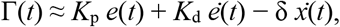

with proportional gain *K*_p_ = κ / *g*′ and derivative gain *K*_d_ = κδ / *g*′. Because the mapping ϕ = *g*(*x*) is linear with constant slope *g*′, the residual term reduces to joint damping 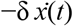, proportional to the plant velocity.

The derivative gain scales linearly with the delay while the proportional gain does not, so the gain ratio reports the delay directly,

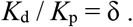

The result is model-independent: it requires only near synchrony, a delay, and an odd coupling function, and holds for any oscillator type. Synchronization is therefore *sufficient* to implement feedback control.

### Simulations

To verify the prediction we drove a second-order plant (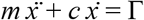, *m* = 1, *c* = 0.8) with sinusoidal coupling Γ = κ sin[ϕ_L_(*t*−δ) − ϕ_F_(*t*)] (κ = 2, δ = 0.12 s), with the lead phase encoding a step target and the follower phase tracking the plant. The coupled-oscillator step response closely matches an ideal PD controller using the analytically predicted gains *K*_p_ = κ, *K*_d_ = κδ (Figure 2, left).

**Figure 2.**
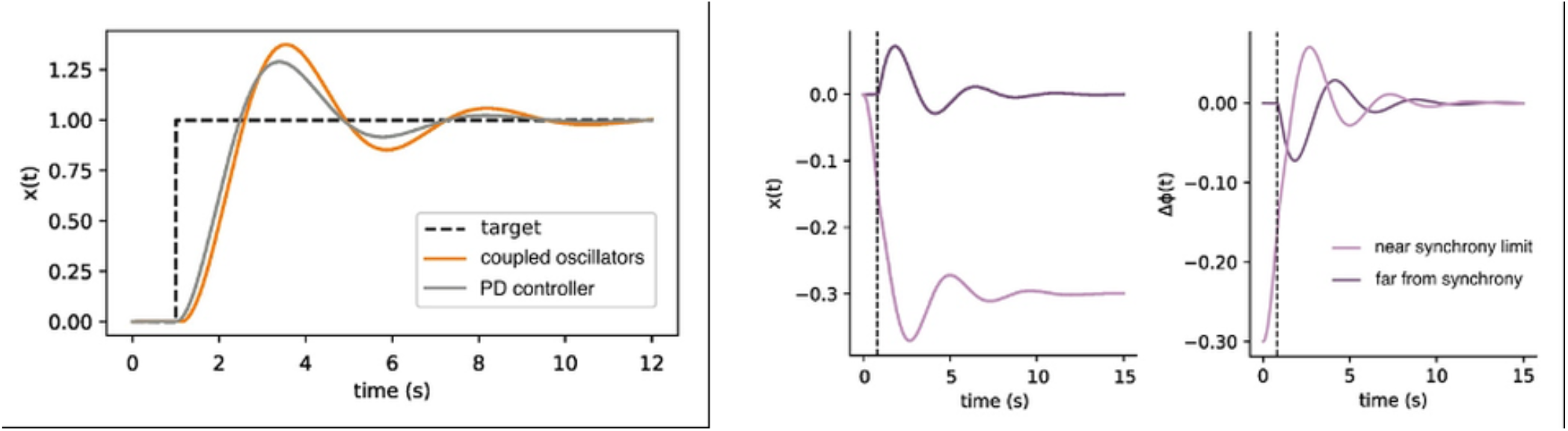
Coupled oscillators implement PD control near synchrony and degrade away from it. Left: step response of the coupled-oscillator plant (κ = 2, δ = 0.12 s) overlaid on an ideal PD controller with matched gains K_p_ = κ = 2, K_d = κδ = 0.24; the two trajectories agree closely. Right: response to a brief perturbation (Δx applied for 0.15 s) within versus outside the near-synchrony limit. The phase difference Δϕ(t) recovers in both cases, but the plant state x(t) settles back to baseline only near synchrony, retaining an offset far from synchrony.

The correspondence holds only near synchrony. Applying a brief perturbation while the oscillators are near synchrony (Δϕ ≈ 0) versus away from it (Δϕ ≈ −0.15 rad) shows that the phase difference recovers in both cases, but the plant state returns to baseline only near synchrony; far from synchrony a sustained offset remains (Figure 2, right). Control quality thus degrades systematically as synchrony is lost, consistent with the linearization breaking down.

### Experimental signatures

A system implementing control via synchronization of two oscillators should exhibit the following signatures: (i) the regulation error should be encoded in the phase error between the coupled populations; (ii) disrupting synchrony should specifically degrade control performance while leaving timing largely intact; and (iii) the effective derivative-to-proportional gain ratio should track the interaction delay, *K*_d_ / *K*_p_ ≈ δ. Importantly, these predictions need not hold for more complex oscillator networks (i.e. more than two oscillators).

## Conclusion

Delayed, odd-function coupling between two near-synchronous oscillators yields an effective PD feedback law plus damping, in the reduced dynamics, independent of oscillator model. Coupled-oscillator synchronization and feedback control, usually treated as separate principles in the motor system, are here two aspects of one mechanism: near synchrony, the synchronization dynamics itself *is* the controller implementation. This reframes pervasive motor oscillations as a plausible substrate for feedback control and offers concrete predictions to test the underlying mechanism in candidate systems.

## Acknowledgements

I am supported by a Lundbeck Foundation Postdoctoral Fellowship (Grant no. R483-2024-1882). I thank Jens Bo Nielsen and Simon Farmer for feedback and Jonathan Scharff Nielsen for proofreading.

